# Age and anatomical region related differences in vascularization of the porcine meniscus using micro-computed tomography imaging

**DOI:** 10.1101/2023.11.08.565772

**Authors:** V-P. Karjalainen, V. R. Herrera M, S. Modina, G. M. Peretti, M. Pallaoro, K. Elkhouly, S. Saarakkala, A. Mobasheri, A. Di Giancamillo, M.A. Finnilä

## Abstract

Meniscal lesions in vascularized regions are known to regenerate while lack of vascular supply leads to poor healing. Here we developed and validated novel methodology for three-dimensional structural analysis of meniscal vascular structures with high-resolution micro-computed tomography (µCT).

We collected porcine medial menisci from 10 neonatal (not-developed meniscus, n-) and 10 adults (fully developed meniscus, a-). The menisci were cut into anatomical regions (anterior horn (n-AH & a-AH), central body (n-CB & a-CB), and posterior horn (n-PH & a-PH).

Specimens were cut in half, fixed, and one specimen underwent critical point drying and µCT imaging, while other specimen underwent immunohistochemistry and vascularity biomarker CD31 staining for validation of µCT. Parameters describing vascular structures were calculated from µCT.

The vascular network in neonatal spread throughout meniscus, while in adult was limited to a few vessels in outer region, mostly on femoral side. a-AH, a-CB, and a-PH had three, five, and seven times greater vascular volume than neonate, respectively. Moreover, thickness of blood vessels, in three regions, was six times higher in adult than in newborn. Finally, a-PH appeared to have thicker blood vessels than both a-AH and a-CB.

For the first time, critical point drying-based µCT imaging allowed detailed three-dimensional visualization and quantitative analysis of vascularized meniscal structures. We showed more vascularity in neonatal menisci, while adult menisci had fewer and thicker vascularity especially limited to the femoral surface which is involved in load transmission response, thus suggesting how nutritional support in this area of the outer zone is more necessary.

## Introduction

The menisci are C-shaped fibrocartilaginous tissues located in the knee joint. They are known to have a primary function in biomechanical properties of the knee joint, like load bearing, joint stability, and shock absorption, that are provided by a complex collagenous extracellular matrix (ECM) organization, including proteoglycans and vascular supply^1–3^. However, menisci are prone for degeneration, usually caused by tearing or osteoarthritis (OA), the spontaneous healing possibilities are weak due to the limited vascular supply of the meniscus^4,5^.

In a developed meniscus “red-red”, “red-white”, and “white-white” – zones can be identified based on their assessment of blood supply with each zone radially covering approximately a third in developed meniscus^6–8^. The peripheral red-red zone is composed of a high vascular supply; the red-white zone in the middle third has limited vascular supply, and finally, the inner third, the white-white zone, is completely avascular. The highly vascularized red-red zone retains the ability to spontaneously heal damage, while in the inner avascular area, the healing potential remains low without pre-emptive surgery. Overall, vascular supply through the meniscal tissue is suggested to have an important role in the healing of the meniscus through revascularization and overall increased healing in the outer red-red regions of the meniscus ^5,9–12^.

Previous studies have reported that vascularity decreases with increasing age in human menisci^7,13–16^. The high degree of vascularity is important for the early growth and maturation of the meniscus, with blood supply covering most of the meniscus width and length^17^. However, in the developed meniscus of adults, only the outer third remains vascularized to some extent in both animals and humans^18^. In addition, regional differences have been found between the horns and body of the menisci with more vascularization found near the horns of the meniscus, that reside further away from the load-bearing areas^1,7,19^.

We have previously shown that human meniscal microstructures can be visualized and quantitatively analyzed with micro-computed tomography (µCT) using drying-based sample processing^20,21^. Critical point drying (CPD)-based drying processing allows to utilize intrinsic contrast of the soft tissue matrix in the µCT imaging. However, the vascular supply of the porcine meniscus has not been quantitatively analyzed in 3D.

In this study: our objectives were to: 1) analyze the vascular supply of ex vivo porcine meniscus samples in 3D using CPD-based µCT imaging protocol in neonatal and adult animals, thus performing validation of µCT imaging with immunostaining of meniscal vascularization, 2) quantitatively compare the vascular supply (vascular fraction percentage) of medial menisci in the anterior horn, central body, and posterior horn of neonatal and adult (fully developed) porcine and subsequently compare the vessel morphology with blood vessel volume, fraction, thickness, and penetration depth and lastly analyze the presence of calcification in the adult menisci. We focused on the medial meniscus because it is more firmly attached to the tibial plateau than the lateral one and for this reason, is more subjected to pathologies.

## Method

### Tissue sample preparation

Medial porcine meniscus specimens from 10 neonates were collected from stillbirth pigs provided by a local breeding farm. We also selected 10 medial porcine meniscus specimens from adult (3-4 years old) pigs from a local slaughterhouse. The neonatal menisci specimens were chosen to represent a group of completely vascularized menisci, or not fully developed neonatal population, while the 3-4-years-old menisci specimens are fully developed and thus were chosen to represent the adult population. The samples were cleaned under a sterile hood in phosphate-buffered saline (PBS, Thermo Fisher Scientific) and fixed for at least 14 days in 10% neutral buffered formalin (Bio-Optica Milano, Milan, Italy). All menisci were radially sectioned into three anatomical regions: anterior horn, central body, and posterior horn.

Hereafter, these neonatal and adult anatomical regions are referred to as n-AH (Neonatal Anterior horn), n-CB (Neonatal Central body),n-PH (Neonatal Posterior horn), and a-AH (Adult Anterior horn), a-CB (Adult Central body), a-PH (Adult Posterior horn), respectively. The specimens were further cut into two few millimeters-thick tissue sections for µCT imaging and histological sample processing. No animals were sacrificed for the purposes of this study: all cadavers were used according to the principles stated by the EC Directive 86/609/EE about the protection of animals used for experimental and other scientific purposes.

### Immunohistological validation of vascularity

Fixed histology specimens were washed for half an hour in running water and decalcified for at least 5 days in a solution consisting of citric (7.5% Sigma-Aldrich) and formic acid (25%, DITTA). The samples were washed again in running water for one and a half hours, in distilled water for 30 minutes, and dehydrated in an increasing scale of ethanol (70% overnight [O.N.], 80% 30 minutes, 95% 60 minutes, 2 x 100% 45 minutes). They were then clarified in xylene (two 45-minute washes) and embedded in paraffin for sectioning. 4µm thick sections were cut and stained with vascular-related markers in immunostaining. The anti-CD31 antibody (Abcam) was used as a vascular marker as described by Canciani et al (2021)^22^. Briefly, sections were deparaffinized in xylene followed by a descending scale of alcohols to water (2 x xylene 5 minutes, 2 x 100% ethanol 5 minutes, 95% ethanol 5 minutes, 70% ethanol 5 minutes, distilled water 5 minutes). Endogenous peroxidase blocking was performed with 10% hydrogen peroxide (H_2_O_2_) for 10 minutes and a 5-minute PBS wash was performed. A pH 6 citrate buffer retrieval for 5 minutes in the microwave followed by half an hour of cooling was performed twice. One hour of blocking with 10% Normal Goat Serum (NGS, Thermo Fisher Scientific) preceded incubation with the antibody diluted 1:50 in PBS + 10% NGS O.N. at +4°C. The following day the slides were washed in PBS for 5 minutes and incubated with Envision anti-rabbit for 3 hours at room temperature (RT). To develop, 3,3’-diaminobenzidine (DAB, Thermo Fisher Scientific) was used (18 minutes). The samples were then counterstained with Meyer’s hematoxylin for two minutes and re-hydrated in an increasing scale of alcohols and clarified in xylene. Finally, Eukitt mounting medium (Sigma-Aldrich) was used to mount the coverslip. B-1000 Optika microscope (OPTIKA) was employed to acquire the images. The described immunohistological method was used as a validation for the µCT method.

### µCT imaging of meniscal vascularization

After fixation, the µCT pieces were dehydrated in an ascending ethanol series (30%-50%-70%-80%-90%-96%-100%) for a minimum of 4 hours in each step. To obtain optimal soft tissue contrast for µCT imaging, samples were dried using a critical point drying (CPD) device (K850, Quorum). Image acquisition was conducted with a desktop µCT system (SkyScan 1272, Bruker microCT, Kontich, Belgium) with the following settings: tube voltage 40 kV; tube current 250 μA; no additional filtering; isotropic voxel size 3.3 μm; the number of projections 2384; averaging 3 frames/projection; and exposure time 1515 ms. Image reconstruction was done with NRecon software (v1.7.1.6. Bruker microCT) with beam-hardening and ring-artifact corrections applied.

### Volumetric 3D analysis of meniscal vascularization

For each sample, cross-sectional µCT images were used for the 3D segmentation of vascularity in the meniscus using Dragonfly (Version 2022.1, Object Research Systems (ORS) Inc, Montreal, Canada). Although the vascularity was not stained with a specific contrast agent, the CPD-based drying process gave it a distinct contrast difference when compared to the surrounding collagenous extracellular matrix. Moreover, distinct morphology and connected network of blood vessels allowed segmentation of the vascularity. Thus, local thresholding together with manual segmentation was used to attain the vascularized volumes. Furthermore, despeckling was used to remove noise. We produced vascular volumes of all samples and measured four descriptive parameters for vascularity: a) volume, b) fraction, c) thickness, and d) depth. In addition, total tissue volume was calculated to clarify the size relationship between the groups. 3D volumetric analysis of the segmented vascularity was performed using CTAn (Bruker MicroCT, version 1.20.3). The penetration depth of blood vessels was calculated as a fraction of the maximum radial depth of the meniscus.

Visualization of all 3D µCT-images was done with CTVox (Bruker MicroCT, version 3.3.0).

### Statistical analyses

Statistical analysis was performed using GraphPad Prism 8 (GraphPad Software, Inc. San Diego, CA). Group-wise comparison of estimated means between anatomical regions of the meniscus was conducted with a Linear Mixed Model in Prism Graphpad (V. 8.4.2) with matching anatomical regions within single meniscus (n-AH vs. n-CB vs n-PH), assumption of sphericity, and with Bonferroni correction. Group-wise comparison of estimated means between age groups of menisci (n-AH vs. a-AH) was done with Geisser-Greenhouse sphericity correction and Bonferroni correction. The descriptive data of menisci are provided in scatterplots with 95% confidence intervals (95% CI). Estimated means and estimated differences between groups are provided in Supplement Tables S1 and S2.

## Results

Immunohistochemistry – validation of vascularity

CD31 immunostaining allowed the identification of blood vessels in the tissue. In the neonatal menisci, blood vessels were identified throughout the tissue, both in the inner zone (Fig. 1A, black circles with blood vessels) and the outer zone (Fig. 1B, black circles with blood vessels); while in the adult menisci, the inner zone appears completely devoid of blood vessels (Fig. 1C) which, however, remain in the outer zone (Fig. 1D, black circles with blood vessels). This difference existed in the three meniscal portions (PH, CB, AH).

**Figure 1.**
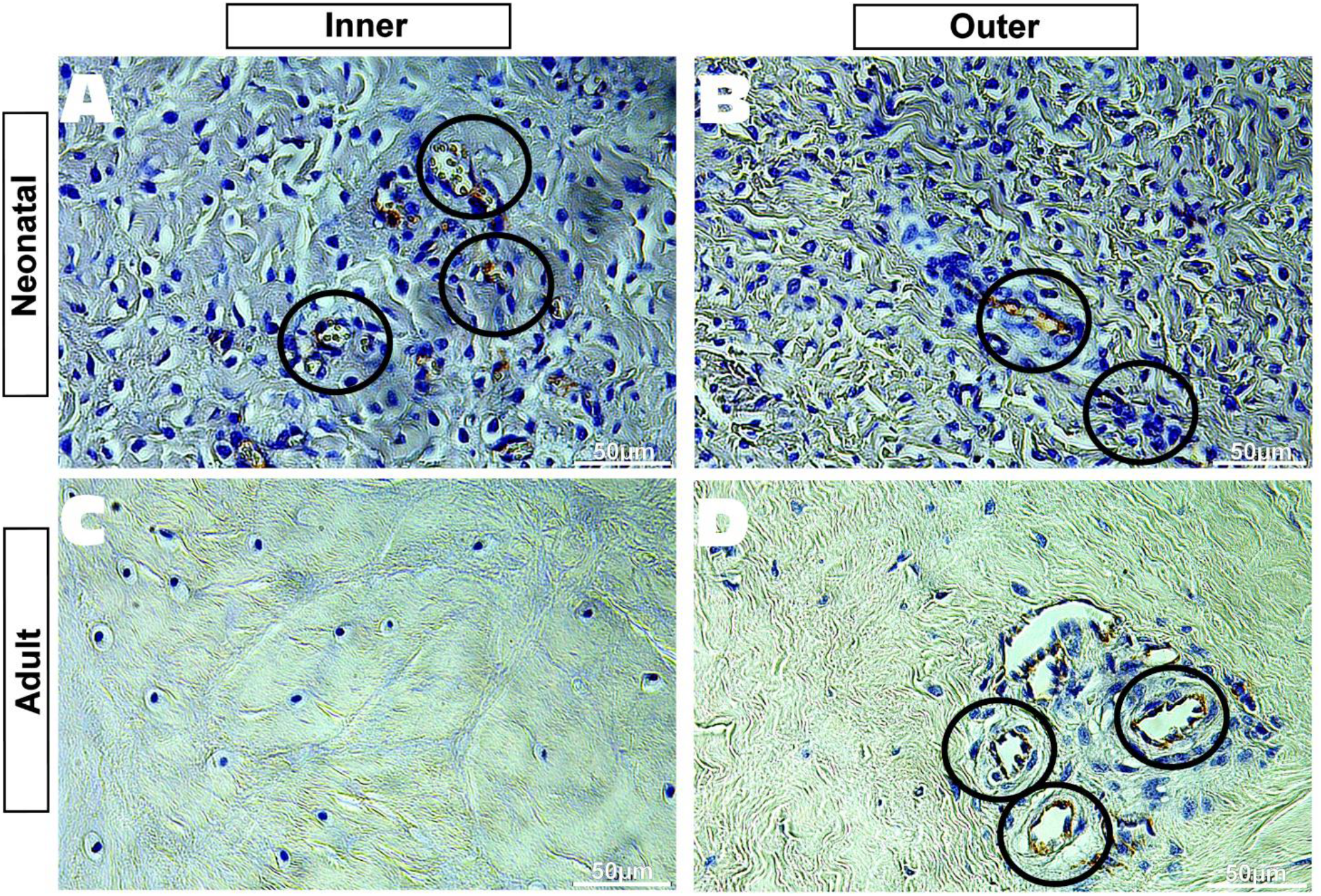
CD31 immunohistochemical staining of newborn (A-B) and adult (C-D) pig meniscus; representative images of the posterior horn. Black circles indicate blood vessels, containing red blood cells. Scale bar: 50 um.

### µCT imaging of meniscal vascularization

The above-presented results of the CD31-stained immunohistology slices were used to validate the µCT segmentations. Representative 2D immunohistology images, 2D µCT slices of adjacent meniscus pieces, and 3D visualization of µCT images are displayed in Fig. 2.

**Figure 2.**
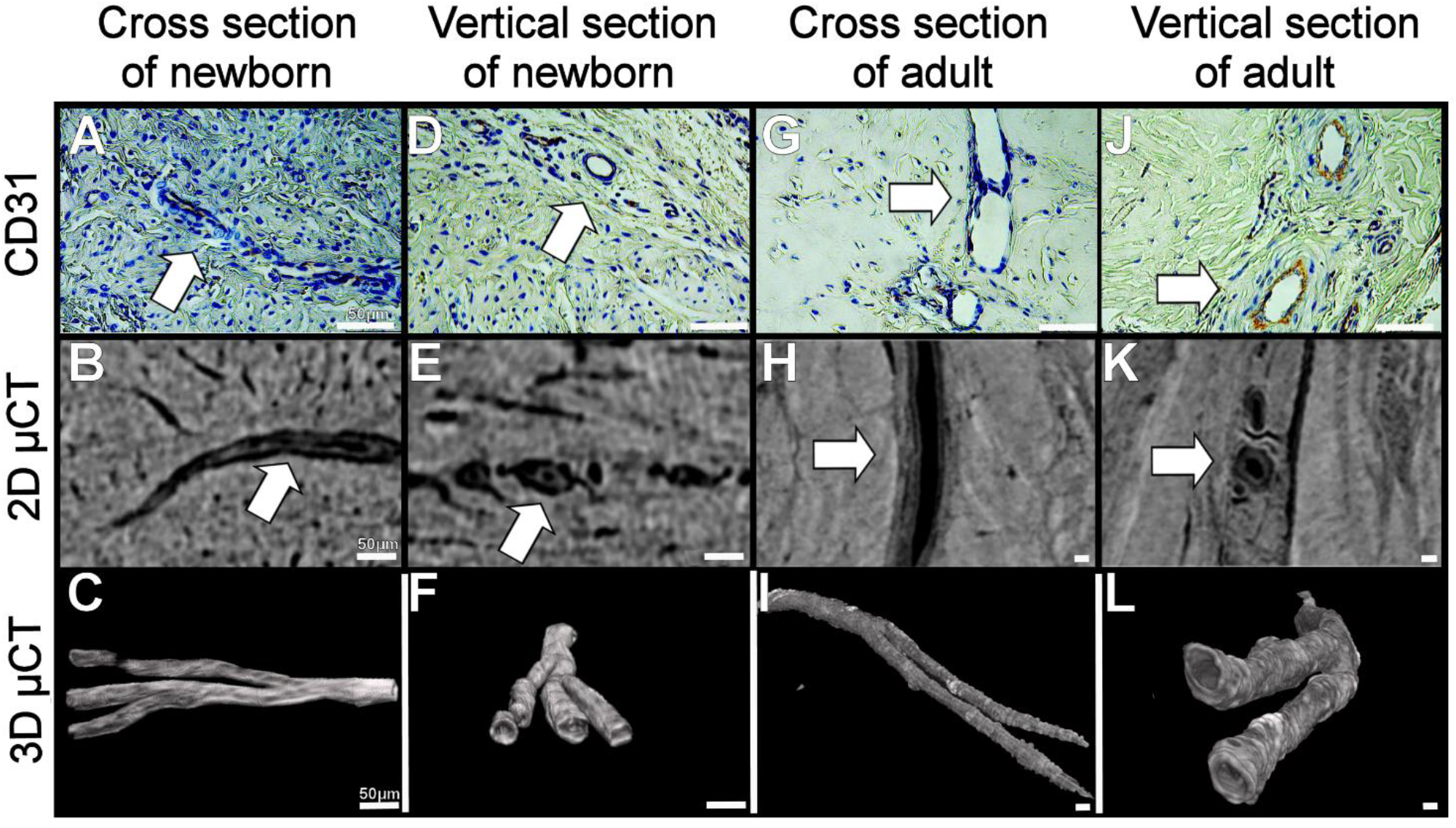
The representative posterior horn of 2D immunohistology image (A/D/G/J), 2D µCT slices of adjacent meniscus piece (B/E/H/K), and 3D visualization of µCT images (C/F/I/L) of both newborn (A-F) and adult (G-L) in vertical (A-C, G-I) and cross-sectional (D-F, J-L) slices. Assessment of blood vessels shows similar morphology and size between µCT-images and CD31-stained histological slices. In addition, the walls and lumen of blood vessels are well visible. Thicker and larger blood vessels of adult menisci can be seen in both histological and µCT-images. Scale bars: 50 µm.

Visual evaluation of both µCT cross-sectional images and CD31 immunohistology showed similar morphology and size in blood vessels between methods (Fig. 2 A vs B, D vs E, G vs H, J vs K, white arrows). The CPD-based drying method seems to preserve the tissue and vascular network intact for accurate µCT imaging. Branching and more complex vascularity networks can only be seen in 3D from µCT visualization compared to standard histology (Fig. 2C/F/I/L).

The vascularization network in CPD-treated meniscal samples was successfully depicted in 3D using µCT imaging. In Figure 3, we show example images of µCT 3D volumes from neonatal and adult porcine menisci, their regions, and their segmented vascularization.

**Figure 3.**
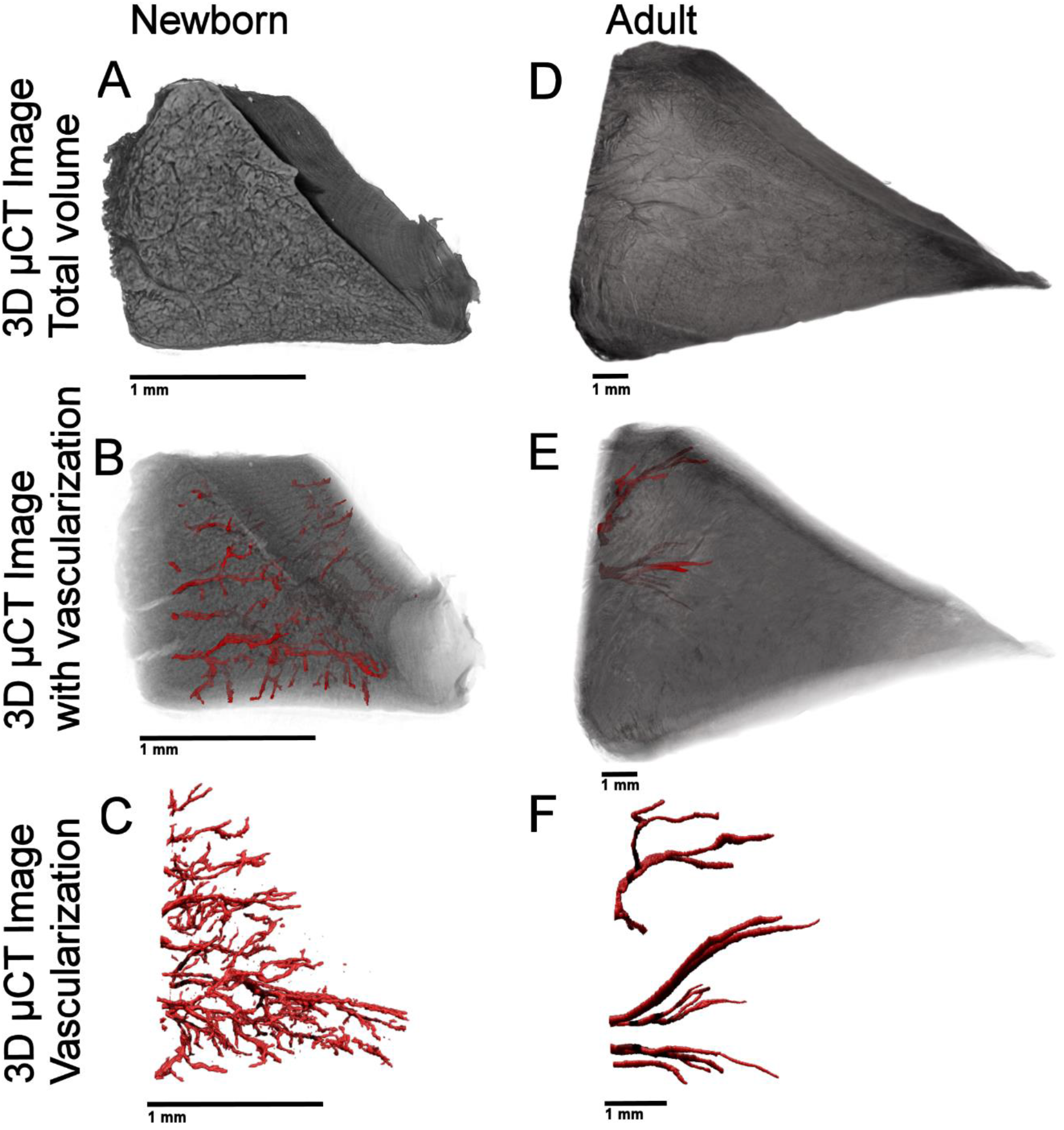
Representative µCT 3D volumes from the posterior horn of neonatal (A-C) and adult groups (D-F). Meniscal vascularization displayed together with soft tissue shows higher vascularity in the neonatal group (B), penetrating throughout the whole meniscus, while the adult group is found only in the outer third, mostly on the femoral side (E, black arrow). The analyzed meniscal vascularization were depicted in newborns (C) and adults (F).

Specifically, Figure 3A shows the 3D scan of the tissue, Figure 3B shows the vessels inside the meniscus, and Figure 3C shows the isolated vascular network, clearly highlighting the differences between the newborn and the adult. The vascular network in the newborn spreads from the outer to the inner region, while in the adult it is limited to a few vessels in the outer region, mostly located on the femoral side (Supplementary videos S1 and S2, Fig. 3E, black arrow).

### Volumetric 3D analysis of meniscal vascularization

Quantitative measurements of the vascular volume, the anterior horn, central body, and posterior horn of the adult were greater than the neonate, respectively (AH: p<0,001, CB: p<0,001 and PH: p<0,001) (Fig. 4A). Within the same age group, no differences were observed between the various portions of the newborn pig, but in the adult, both the anterior horn and the central body were significantly different from the posterior horn (p<0,001) (Fig.4A).

**Figure 4.**
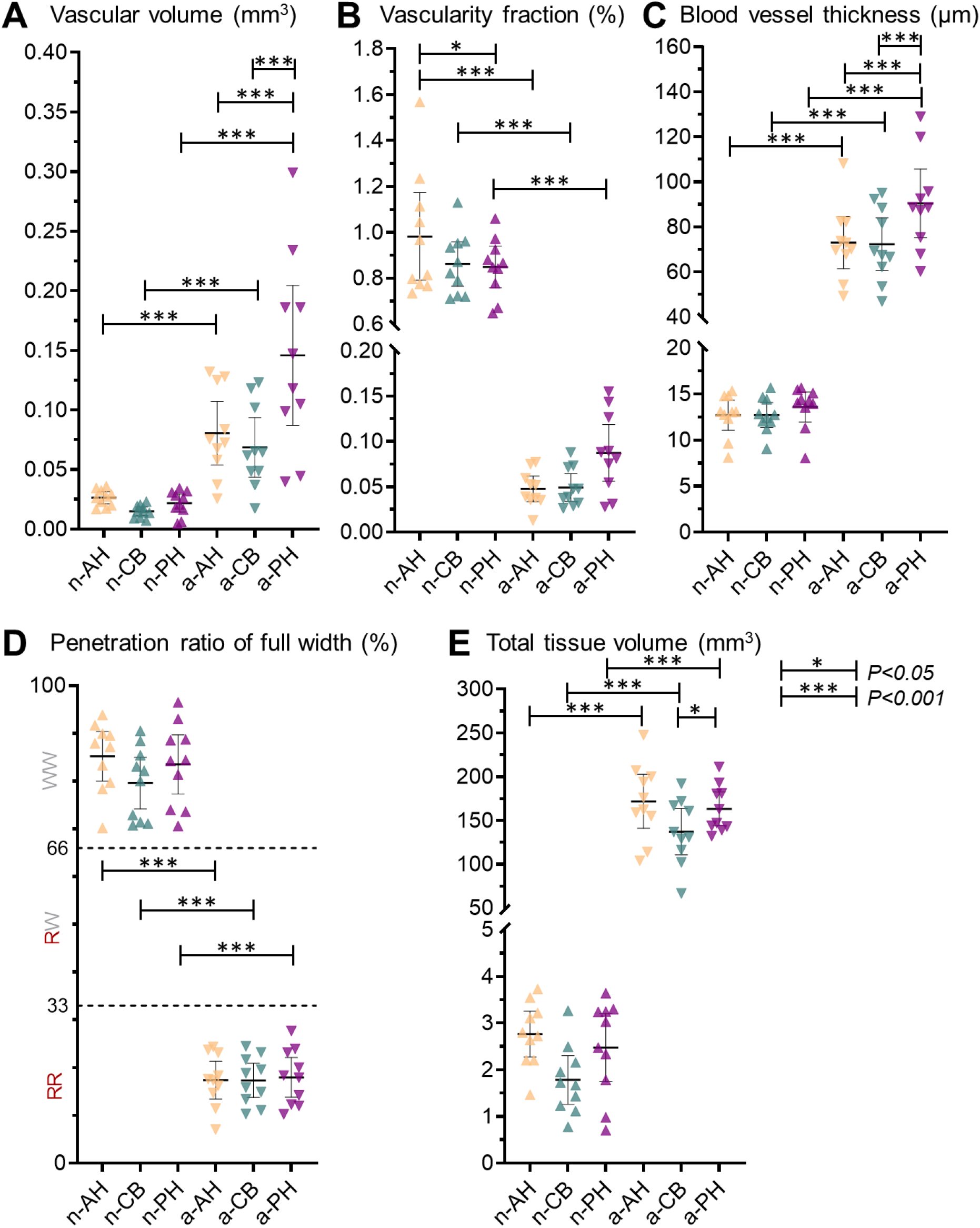
A) Vascular volume comparison B) Vascular fraction comparison of tissue fraction. C) Blood vessel thickness. D) Penetration depth of blood vessels in ratio of radial tissue length and expected regions (RR = red-red, RW = red-white, WW = white-white) of vascularity. E) Total tissue volume of the meniscus to clarify the size relationship between the groups. Comparisons were done between newborn and adult meniscus in their anatomical portions (AH= anterior horn, CB= central body, PH= posterior horn). Group-wise comparison of estimated means between anatomical regions of the meniscus was conducted with a Linear Mixed Model in Prism Graphpad (V. 8.4.2) with matching anatomical regions within a single meniscus (n-AH vs. n-CB vs. n-PH). Group-wise comparison of estimated means was done between age groups of menisci (for example, n-AH vs. a-AH) N= 10, data are expressed as mean ± 95% C.I. ***= p<0,001.

In the three portions (AH, CB, PH) a significant reduction (p<0,001, Fig. 4B) was observed in the adult vascular fraction compared to newborn animals. n-AH appeared to have a higher vascular fraction compared to n-PH (p<0,05, Fig. 4B), but no differences were observed comparing n-CB and n-PH. There were no statistically significant differences between the three portions in the adult animal (p>0,05, Fig. 4B).

Concerning the thickness of blood vessels, in the three portions (AH, CB, PH) the adult had an average of about six times higher than the newborn (p<0,001, Fig. 4C). In Figure 4C, we found that the average blood vessel thickness in neonatal menisci was highest in n-PH group with n-AH and n-CB having a similar average thickness (p>0,05, Fig. 4C). Furthermore, a similar trend was found in adult menisci with highest blood vessel thickness found in a-PH group (p<0,001, Fig. 4C), while same thickness was perceived between a-AH and a-CB (p>0,05, Fig. 4C).

Regarding the depth of penetration of the vessels into the tissue, it was drastically reduced in the transition from the newborn to the adult animal in the three sections (p<0,001, Fig. 4D), but no significant differences between them neither in the newborn nor in the adult (Fig. 4D). Numerical results are found in Supplement Table I with estimated differences between groups in Supplement Table II.

### Calcifications

Incidentally, we found calcified clusters in three samples of the a-AH group. In Figure 5, we present the scatterplot of vascular fraction between calcified and non-calcified tissue. After splitting the a-AH group, we see that the calcified group (n=3) had a seemingly higher vascular fraction when compared to non-calcified anterior horns (n=7). However, we did not perform any statistical analysis for these as the number of samples in the calcified group was low.

**Figure 5.**
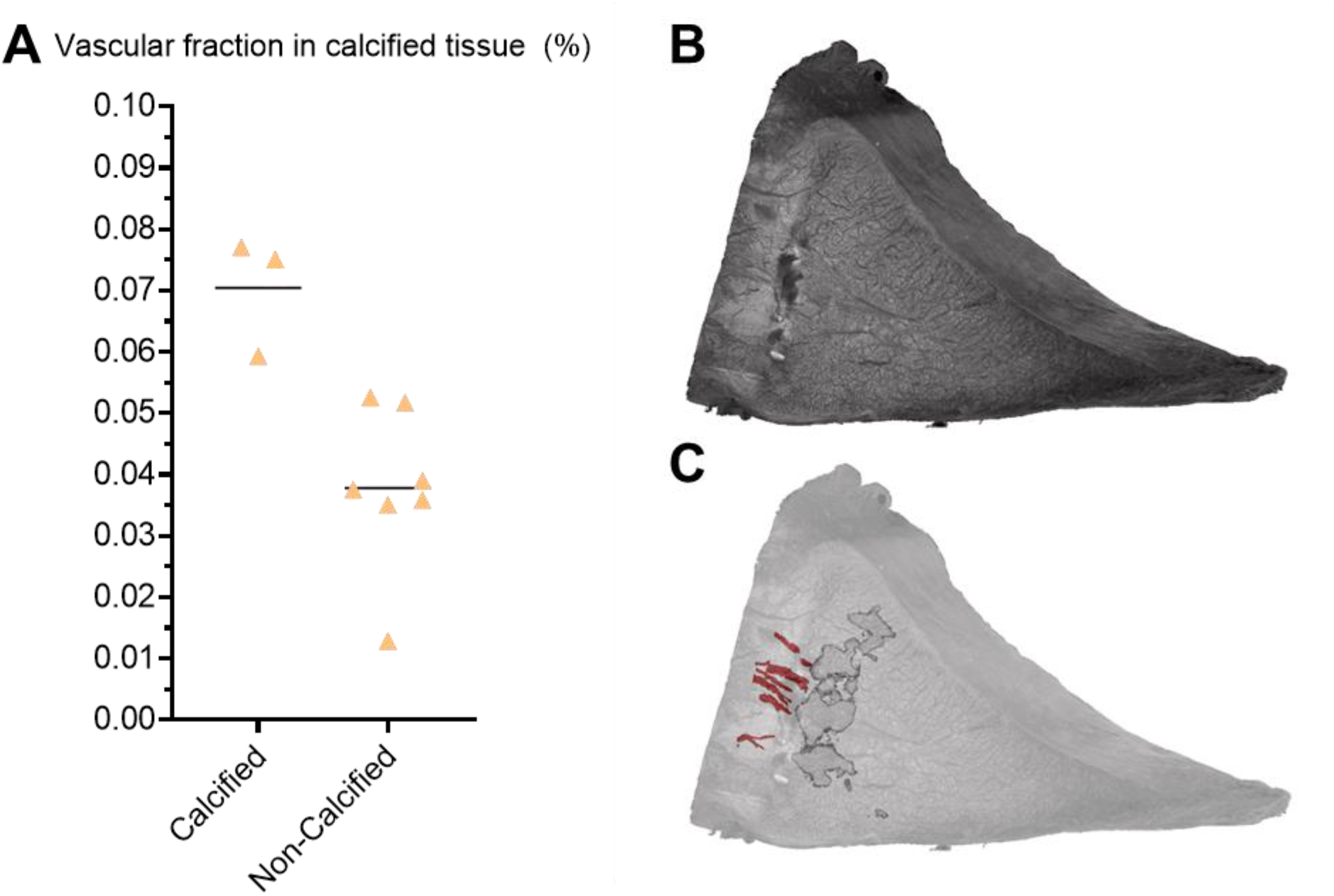
A) Mean vascular fraction (%) in the anterior horn of adult porcine meniscus (a-AH) between calcified and non-calcified tissue. Calcified menisci had a higher average vascular fraction denoting higher vascularity overall compared to non-calcified menisci. B) 3D µCT image of the medial anterior horn of adult porcine meniscus with calcification clusters inside the tissue C) 3D µCT image showing calcified structures together with vascularization in the meniscus.

## Discussion

CPD-based sample drying protocol together with high-resolution µCT imaging is a feasible method to visualize and quantitatively analyze the vascular network of porcine meniscus in 3D. The results of volumetric analyses indicate not only substantial differences in vascular supply between neonatal and adult groups, but also regional differences between anterior horn, central body, and posterior horn. Decreased vascular content in adult groups confirms a strong association between age and vascularity of meniscus already observed in our previous research in pigs’ developmental studies ^23–25^. Furthermore, vascularity seems to decrease the most in the anterior horn and least in the posterior horn of the meniscus, suggesting regional differences. To our knowledge, this is the first study to quantitatively study the vascularization in 3D between neonatal and adult porcine menisci considering the three zones.

CPD-based µCT imaging protocol was validated using CD31-stained immunohistology. CPD treatment seemingly preserved the tissue when compared to the immunohistology with blood vessels being visible in both immunohistology and µCT images while sharing similar morphology and size. Qualitatively, both immunostaining and µCT showed fully vascularized neonatal menisci while vascularity in adult menisci seemed to only contain vessels in the outer region. Branching and more complex vascularity networks can be seen in 3D from µCT visualization. Importantly, branching and 3D structures can only be detected using µCT-imaging compared to the standard 2D immunohistology. However, immunohistology is an important complementary method for µCT in identifying tissue structures, that do not necessarily have different attenuating properties in µCT images.

From our volumetric µCT analyses, we found a higher vascularity volume fraction in the neonatal menisci when compared to adult menisci, suggesting a decrease in vascular volume fraction with age. Qualitatively, the neonatal menisci showed greater branching and connectivity compared to adult menisci. Our findings are supported by previous cadaver studies measuring vascular density, where overall vascular density decreased with increasing age in human menisci ^14,15,26,27^. In our study, the neonatal menisci were seemingly less wedge shaped and relatively thicker radially compared to the more normal, wedge-shaped adult menisci. Supporting findings have been found by the non-developed neonatal meniscus being reported to be more heightened surfaces compared to the wedge shape of developed adult meniscus^26^. Furthermore, in our previous study, we reported that age-related angiogenesis factors are highly expressed in neonatal porcine meniscus^23^. The reason for a sharp decrease in vascularity after birth has been thought to be derived from the start of biomechanical stress and loading in the meniscus after birth, especially in the inner white zones causing extrusion in the meniscus^26^. Nutrition via diffusion through a large vascularization network in the meniscus is important before the loading starts around 2 years of age in humans, after which the nutrition of these areas occurs via diffusion^26^. If meniscus tears or degenerates, nutrition is required for healing, possibly leading to revascularization in the form of increased vascular volume and blood flow as a response to injury^28^. In OA, it has been identified to promote angiogenesis and increase vascularity in the knee joint and meniscus^29–31^. Moreover, meniscal lesions have also been reported to cause inflammation and swelling in the OA tissue^32^. Thus, the revascularization to heal tears in avascular areas in healthier and compressed tissue may be suppressed by tightly organized collagenous ECM, while revascularization could manifest more easily in more degenerated, sparser tissue where reparative cells can infiltrate the tissue^33^. Similarly, revascularization is essential in successful meniscal allograft transplantations. Studying the effect of revascularization in allografts with µCT could be a beneficial 3D method in the future. Analyzing more specifically the meniscus regions, it was observed that in both adult groups, the central body and anterior horn had the lowest volume of blood vessels, which correlates with other studies showing the central body of the meniscus having seemingly less vascularity compared to the horns^16^.

In both neonatal and adult groups, the central body had also the lowest volume and fraction of blood vessels, showing the central body of the meniscus having seemingly less vascularity compared to the horns. The higher vascularity near the horns of the meniscus can be expected as major arteries reside closer to the horns in the periphery capsule of meniscus, supplying the blood vessels^1,7,8,34,35^. Anatomically, the limited blood supply originates mainly from the medial and lateral geniculate arteries: the radial branches, which enter the meniscus at intervals, mainly supply the anterior horn and the posterior horn^36,37^. However, in our study, the largest decrease in vascular content was found in the anterior horn between neonatal and adult groups, while the posterior horn retained the most vascularity and blood vessel thickness with age. The posterior horn inhabiting most vascular content is supported by our previous study showing that posterior horn of adult meniscus regions corresponded to the highest vascular cell activity^38^. It has been studied that in the medial meniscus of humans, the anterior horn is subjected to higher stresses in gait and shows greater stiffness compared to the posterior horn^39–42^. This suggests that lower stress and compressive forces in the porcine gait cycle are localized to the posterior horn, leaving space for thicker and larger vascular organization as per our findings.

We found that blood vessels radially penetrated through most neonatal menisci while in adult menisci they covered less than a third of meniscal tissue, reaching only the outer red-red zone. In our previous angiogenic expression study, we reported that vascular endothelial growth factor (VEGF), which is a specific mitogen for vascular endothelial cells, is evenly expressed between inner, middle, and outer neonatal porcine menisci regions, but significantly increased from inner to the outer part of adult menisci^23^. Correlating results with VEGF expression and our CD31 vascular stain suggest that using µCT imaging can produce important 3D information for the development of degenerative changes or the healing of meniscal transplants^43,44^. Our results about the penetration depth of blood vessels in the adult specimens correlate with that of adult humans, with vessels entering and penetrating only the outer meniscal third or quarter^26^. In our study, the neonatal menisci shows a similar pattern of vascularity as in the meniscus of a newborn human in a previous study^26^. In addition, biomechanical studies have reported similarities in porcine and human meniscus tensile and compression properties and permeability^42,45,46^. Therefore, the porcine meniscus shared similar biomechanical and vascular properties to human meniscus and could be used as a suitable model for fast translation from the pig to the human.

Interestingly, in adult pigs, the vascular network is limited to a few vessels in the outer region, mostly located on the femoral side. In addition, the largest blood vessels in the adult pigs are shown to be located inside the largest radial tie fibers. Arnoczky et al observed that a vascular network extends for a short distance on both the femoral and tibial surfaces of the menisci, but does not contribute to nourishing the meniscal stroma^6^. The same authors found similar vascularization patterns in the two horns. In a previous study of our group, we found that the femoral surface of adult pig meniscus is characterized by a higher quantity of GAGs with the interposition of radial and oblique fibers^47^. These features are responsible for a higher resistance to compressive forces like that acted on by the femoral condyles. On the contrary, the tibial surface shows a circumferential arrangement of the fibers and a poorer GAG presence and cellular spread; these characteristics seem to allow a higher resistance of the tibial surface to traction forces^47^. These peculiar characteristics of the matrix in the femoral side associated with the higher vascular network observed in the same tissue area can be indicative of how the nutritional support linked to the greater biomechanical forces acting on this side of the outer zone is more necessary.

As an incidental finding, we found clusters of calcifications in three anterior horns of adult pigs, two of which were in the red-red zone near the blood vessels and one in the inner white-white region of the meniscus. In early OA, increased expression of VEGF and angiogenesis promotes vascular content leading to ossification in the articular cartilage^29,48^. Moreover, in our previous study, we found that calcifications in the meniscus increase with increasing histopathological degeneration^49^. Our findings show that all three samples inhabiting calcifications had also the highest ratio of the vascular fraction of the anterior horn in adults, suggesting correlation a between meniscus angiogenesis and ossification in the medial anterior horn. We have previously shown that with µCT, you can quantitatively study the calcified tissue in the meniscus in 3D^49^. Thus, the presented method of µCT imaging enables imaging of hard tissues like calcification together with soft tissues like vascularization and could be used to study the effects of ossification and angiogenesis in the meniscus in the future.

Adult animals used in our studies are heavy pigs (<250kg), as they are produced for food production. High body mass index (BMI) and age have been studied to have a significant role in the risk of meniscus injury and OA in both humans and animals^50–54^. Furthermore, our findings of possible revascularization together with ossification in the anterior horn of the medial meniscus of adult pigs suggest a manifestation of early-stage OA. Thus, following the 3Rs, developing a combined early OA and obesity research model based on overweight pigs used in food production could prove a beneficial and sustainable research model option for future studies.

The present study has some important limitations. Firstly, the CPD-based sample drying method preserves the tissue morphology and enables µCT imaging but does cause some shrinking and may not perfectly reflect the sample *in vivo*^55^. Secondly, the incidental calcifications in the menisci can cause streaking artifacts to the image due to differences in soft and hard tissue attenuation, decreasing image quality. Furthermore, the calcified menisci have seemingly higher vascular content compared to non-calcified tissue in the same group, affecting the a-AH group average in vascular volume and fraction measurements. However, in OA, angiogenesis has been suggested to promote endochondral ossification in porcine cartilage, thus spontaneous calcifications in the porcine meniscus could be expected^56^.

In conclusion, µCT imaging allows detailed quantitative volumetric analysis of vascularized meniscal structures in porcine neonatal and adult menisci. In the future, this method could be used to understand the meniscus regeneration and degeneration concerning vascular anatomy in the context of vascular diseases, hypertension, healing of meniscal transplants, and arthritis, or simply for the clinical study of the aged meniscus, which is often accompanied by increased vascularization. In conclusion, µCT imaging allows detailed quantitative volumetric analysis of vascularized meniscal structures.

## Supporting information

Manuscript Supplement

## Acknowledgments

VPK has received a grant from Finnish Cultural Foundation (SKR). MF would like to acknowledge funding from Jane and Aatos Erkko Foundation and Academy of Finland (347445, 353755).

The authors further acknowledge the help of Dragonfly – software (Object Research Systems (ORS) Inc.) in the use of image segmentations.

