## Supplementary material for "Age and anatomical region related differences in vascularization of the porcine meniscus using micro-computed tomography imaging": Manuscript Supplement

**Supplements**

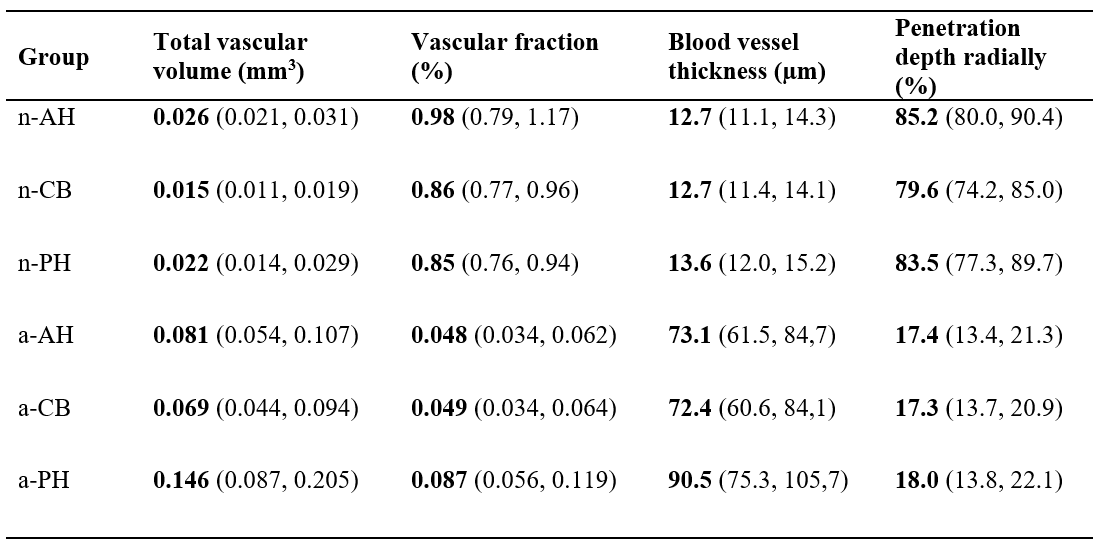

**Supplement Table I.** Mean results of calculated parameters with 95% confidence intervals.

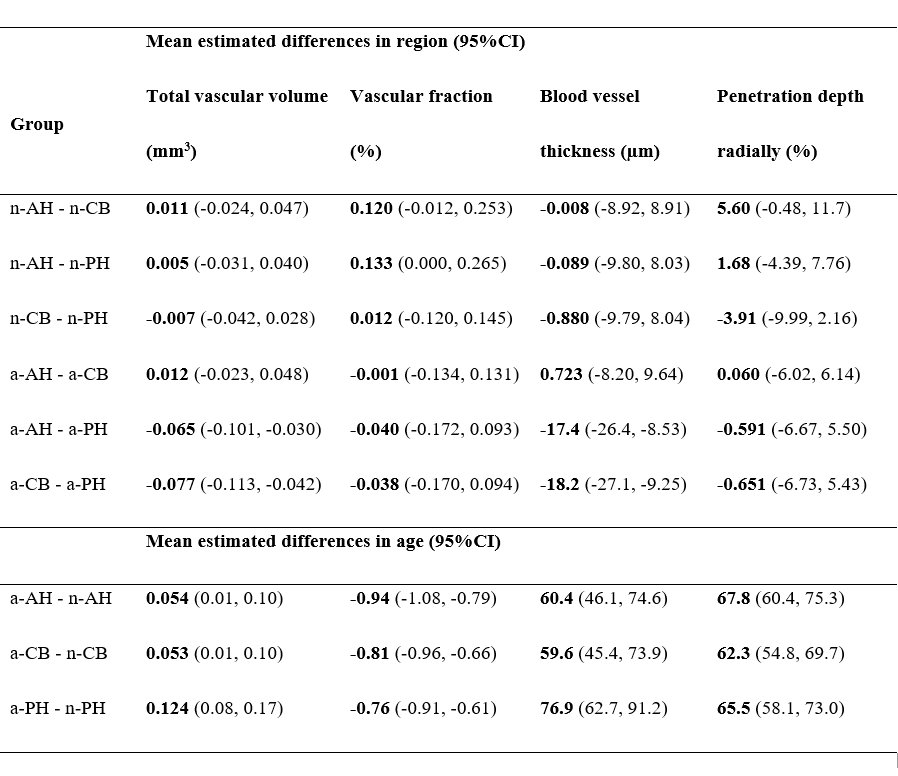

**Supplement Table II.** The mean estimated differences of total vascular volume (mm^3^), vascular fraction (vascular volume / tissue volume, (%)), blood vessel thickness (µm) and penetration depth radially of full tissue width (%) within the following groups: n-AH, n-CB, n-pH, a-AH, a-CB, a-PH. The results are presented as mean with 95% confidence intervals (95%CI).
